# Intrinsic elasticity of nucleosomes is encoded by histone variants and calibrated by their binding partners

**DOI:** 10.1101/392787

**Authors:** Daniël P. Melters, Mary Pitman, Tatini Rakshit, Emilios K Dimitriadis, Minh Bui, Garegin A Papoian, Yamini Dalal

## Abstract

Histone variants fine-tune transcription, replication, DNA damage repair, and faithful chromosome segregation. Whether and how nucleosome variants encode unique mechanical properties to their cognate chromatin structures remains elusive. Here, using novel *in silico* and *in vitro* nanoindentation methods, extending to *in vivo* dissections, we report that histone variant nucleosomes are intrinsically more elastic than their canonical counterparts. Furthermore, binding proteins which discriminate between histone variant nucleosomes suppress this innate elasticity and also compact chromatin. Interestingly, when we overexpress the binding proteins *in vivo*, we also observe increased compaction of chromatin enriched for histone variant nucleosomes, correlating with diminished access. Together, these data suggest a plausible link between innate mechanical properties possessed by histone variant nucleosomes, the adaptability of chromatin states *in vivo*, and the epigenetic plasticity of the underlying locus.

**Significance:** Nucleosomes are the base unit which organize eukaryotic genomes. Besides the canonical histone, histone variants create unique local chromatin domains that fine-tune transcription, replication, DNA damage repair, and faithful chromosome segregation. We developed computational and single-molecule nanoindentation tools to determine mechanical properties of histone variant nucleosomes. We found that the CENP-A nucleosome variant is more elastic than the canonical H3 nucleosome but becomes stiffer when bound to its partner CENP-C. In addition, CENP-C induces cross-array clustering, creating a chromatin state that less accessible. These data suggest that innate material properties of nucleosomes can influence the ultimate chromatin state, thereby influence biological outcomes.

## Introduction

The adaptive nature of chromatin allows a cell to replicate, divide, differentiate, regulate transcription, and repair damaged DNA. In part, the chromatin landscape is shaped by removing old and incorporating new nucleosomes with specific histone variants, and by incorporating covalent modifications (1–8). How different histone variants convey the unique mechanical properties of their nucleosomes to the chromatin fiber, and whether non-canonical nucleosomes modulate chromatin dynamics is a subject of intense study. In contrast to the previous view that chromatin was a mostly static packaging polymer, several recent studies have unveiled a rich conformational landscape of nucleosomes (2). These works raise the intriguing possibility that mechanical properties embedded within evolutionarily distinct nucleosome types might lead to different structural outcomes for the chromatin fiber. Paradoxically, the most evolutionarily divergent histone variant is CENP-A, which is functionally essential across most eukaryotes (9). Another major paradox is that despite being buried in pericentric heterochromatin (10–12), CENP-A chromatin transcriptionally active in most species, suggesting this chromatin is accessible even when bound to kinetochore proteins (13, 14). This puzzling dichotomy can be explained either by intrinsic mechanical properties, or by epigenetic alterations driven by chromatin effectors.

To investigate this salient problem, we developed novel *in silico* and *in vitro* tools to dissect innate mechanical properties of CENP-A nucleosomes relative to their canonical counterparts, in the presence or absence of CENP-A binding partners and extended these findings *in vivo*. We report that the smallest unit of the chromatin fiber can have profound effects on the three dimensional folded properties of chromatin, with implications for the accessibility of that chromatin to the transcriptional machinery.

## Results

### CENP-C^CM^ increases the Young’s modulus of CENP-A *in silico*

We first examined elasticity as a mechanical feature of nucleoprotein complexes, which has never been reported before. Using all-atom molecular dynamics, we measured nucleosome stiffness and examined spontaneous structural distortions that occur in the presence of CENP-C. We ran three simulations for this study: (1) the CENP-A nucleosome core particle (NCP), (2) the CENP-A NCP with one bound rat CENP-C motif of CENP-C (CENP-C^CM^), and (3) the CENP-A NCP with two copies of CENP-C^CM^. As a control, we compared these systems to canonical nucleosomes, H3 (15).

Using these all-atom data, we next developed a novel analytical technique to quantify the elasticity of nucleosomes *in silico*. Briefly, this technique connects structural fluctuations observed in unbiased molecular dynamics simulations, with the nucleosome’s mechanical response, ultimately producing the absolute value of the Young’s modulus (*Methods*). To analyze all-atom simulation data in such a way, we modeled the nucleosomes as mechanically homogenous elastic cylinders vibrating in a thermal bath and calculated the dimensions and fluctuations of these “minimal” cylinders during each simulation trajectory (Figure 1*A*). These analyses predict that the Young’s modulus of CENP-A is far more elastic (6.2 MPa) than that of H3 (9.8 MPa). Interestingly, upon binding either one CENP-C^CM^, or two CENP-C^CM^ fragments (Figure 1*B*), CENP-A nucleosomes adopted a remarkably stiffer configuration (8.2 MPa and 8.7MPa, respectively) much closer to that of H3.

**Figure 1.**
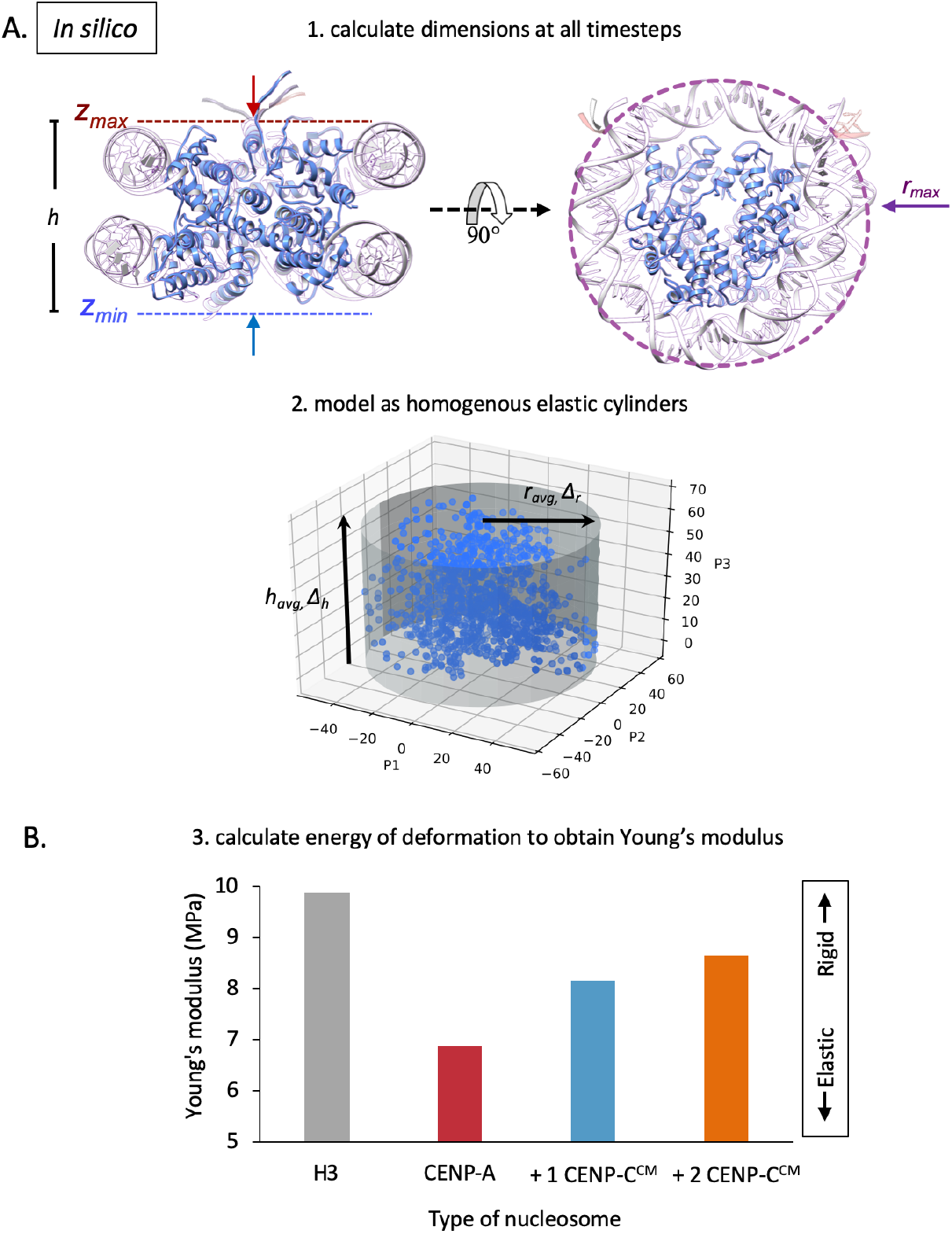
*In silico* analysis predicts that CENP-A nucleosomes are more elastic than H3 nucleosomes. (*A*) To obtain Young’s modulus values from simulation, we measured the *in silico* dimensions of nucleosomes by compression of an encapsulating cylinder programmed to stop at stiffer surfaces resistant to collapse. From the heights, h = z_max_ – z_min_, and the radii, r_max_, of the resulting minimal cylinders we then calculated the average and change in height (h_avg,_ Δh), and radius (r_avg,_ Δr) of each system. (*B*) We treated the nucleosomes as elastic homogenous cylinders, calculated the energy of deformation, and retrieved the Young’s modulus of a cylinder vibrating at equilibrium in a thermal bath.

### CENP-C interactions suppress spontaneous structural distortions of CENP-A nucleosomes

This dramatic alteration in elasticity made us curious to examine conformational changes of CENP-A mononucleosomes that might be induced by CENP-C^CM^. Excitingly, the data revealed a dampening of histone motions relative to each other upon binding of CENP-C^CM^ (*SI Appendix*, Fig. S1*A*). We were curious to assess how these changes would propagate through the DNA. Thus, we investigated DNA gyre sliding and gaping of nucleic acids through *in silico* labeling (*SI Appendix,* Fig. S1*B*). Indeed, a single CENP-C^CM^ fragment dampens CENP-A nucleosome gyre gaping and DNA slides asymmetrically away from the CENP-C^CM^ bound-face of CENP-A nucleosomes (*SI Appendix*, Fig. S1*B*). We performed additional structural analysis to demonstrate local structural flexibility. Altogether, detailed analyses of CENP-A mononucleosomes motions revealed a global dampening of innate motions upon CENP-C^CM^ binding (*SI Appendix*, Fig. S1*A*). On the residue scale, we found that CENP-C^CM^ suppresses residue fluctuations with symmetry breaking in the presence of one fragment (*SI Appendix*, Fig S1*C*). These computational data are in agreement with experimental observations made by sm-FRET and hydrogen/deuterium-exchange mass-spectrometry (16–18) for the CENP-A nucleosome bound to the central domain region of human CENP-C (CENP-C^CD^). The CENP-C^CM^ and CENP-C^CD^ bind to CENP-A nucleosomes through the same mechanisms (19), likely because both domains contain the H2A/H2B acid patch binding motif (RR(S/T)nR) and the CENP-A C-terminal tail binding residues (WW/YW), which are separated by seven residues. Importantly, these two motifs in CENP-C are conserved across plant, fungi, and animal kingdoms (*SI Appendix*, Fig. S2). These data predict that CENP-C dampens motions of CENP-A nucleosomes, and as a consequence, alters mechanical properties of the CENP-A nucleosome.

### CENP-A nucleosomes are more elastic than H3 nucleosomes *in vitro*

To experimentally test this prediction *in vitro*, we turned to nanomechanical force spectroscopy (20, 21). This single-molecule method is used to physically compress and release complexes to directly quantify their elasticity on a nanoscale (22–28). We were surprised to discover that the elasticity of nucleosomes has never been quantified. Therefore, we developed a protocol to perform in-buffer, single-molecule nanoindentation force spectroscopy of nucleosomes (*Methods*).

Using traditional salt dialysis protocols (29, 30), we reconstituted H3 and CENP-A mononucleosomes on linear 187-bp DNA fragments (Fig. 2), or H3 and CENP-A nucleosome arrays on 3 kbp plasmids (Fig. 3*A*). To assess the quality of our reconstitutions, we determine nucleosomal dimensions by AFM, as well as protection from nuclease (MNase) digestion. Consistent with previous work (31, 32), in fluid, *in vitro* reconstituted CENP-A nucleosomes possess dimensions similar to H3 nucleosomes (3.8±0.3 and 3.7±0.3 nm, respectively) (Table 1, *SI Appendix*, Table S1). Likewise, nucleosome arrays yield classical nucleosomal ladders when challenged by MNase (*SI Appendix*, Fig. S3). Using mononucleosomes, we also established nucleosomal orientation, finding that nucleosomes almost always lay flat on mica (Fig. 2).

**Figure 2.**
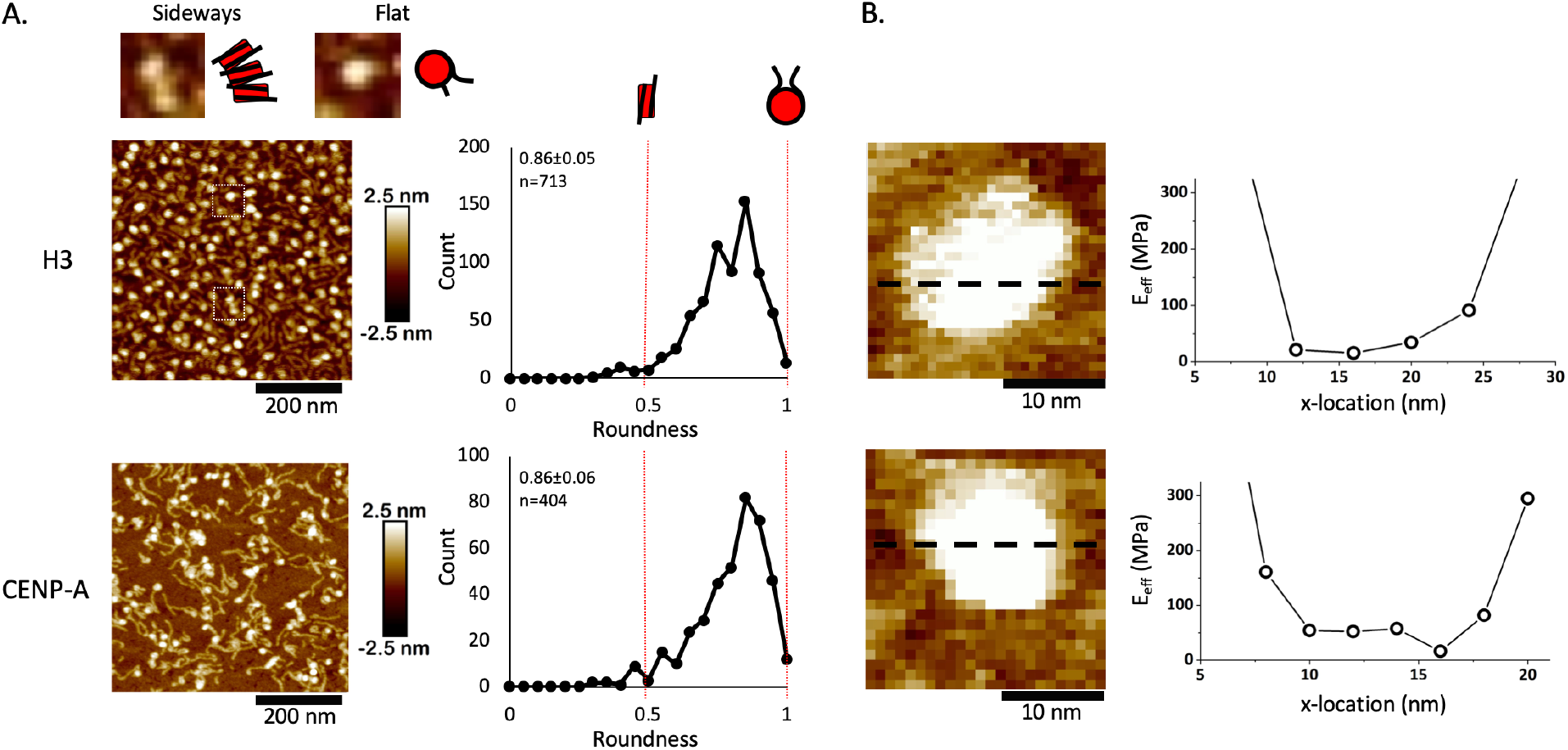
Mononucleosomes lay flat and are uniformly elastic. (*A*) Roundness was measured of either H3 or CENP-A mononucleosomes. A value of 1 would indicate that a nucleosome particle lays flat on the mica surface, whereas a value of 0.5 would indicate a nucleosome particle laying on its side. Almost all nucleosomal particles lay flat. (*B*) Young’s modulus was measured across H3 or CENP-A mononucleosomes to assess whether a nucleosome particle is uniformly elastic. No significant difference in Young’s moduli was observed across either nucleosome.

**Figure 3.**
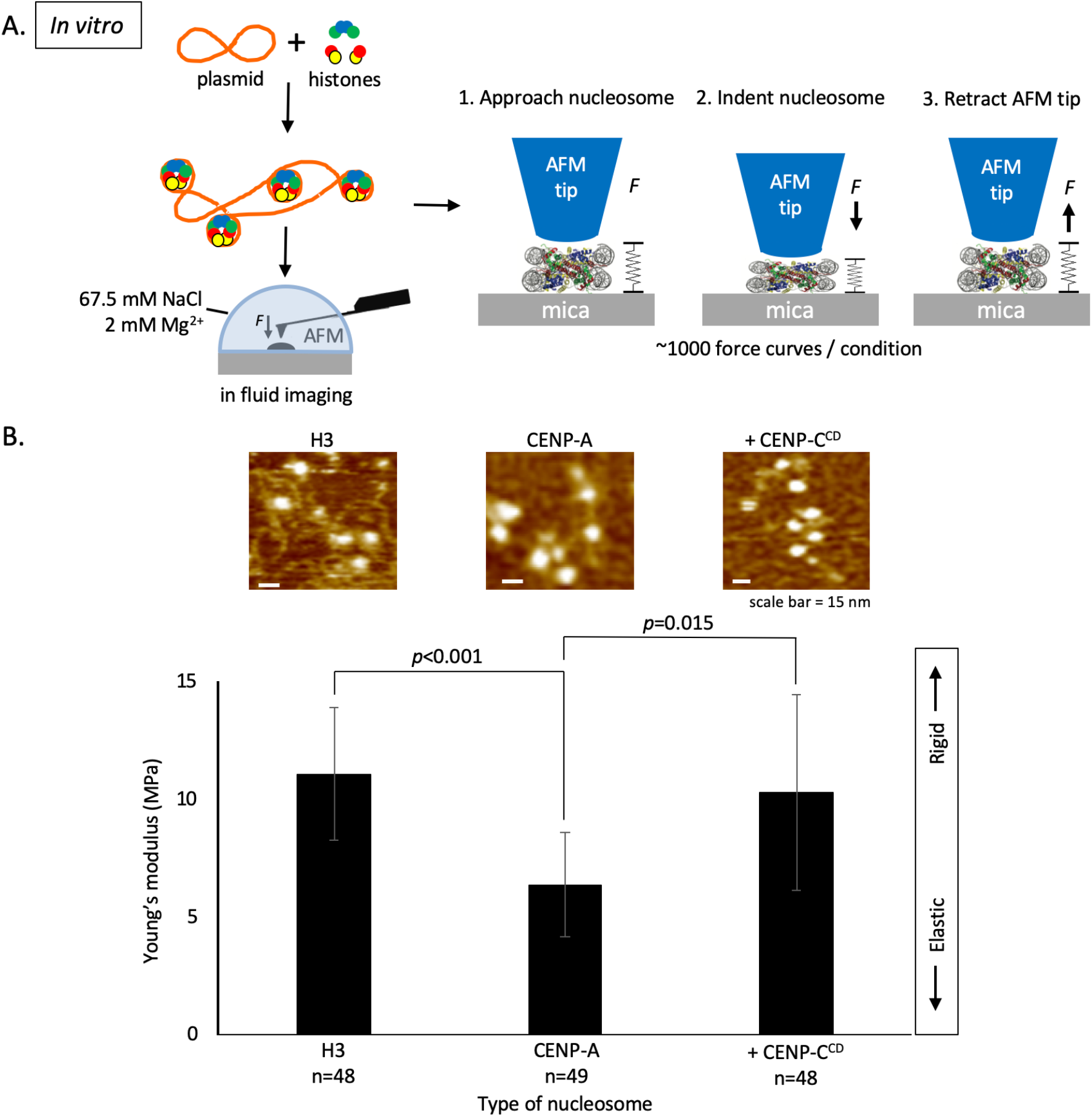
*In vitro* CENP-C^CD^ binding stiffens elastic CENP-A nucleosomes. (*A*) To determine the Young’s modulus of CENP-A and H3 nucleosome arrays, we *in vitro* reconstituted H3 and CENP-A nucleosome arrays by salt dialysis, followed by nanoindentation force spectroscopy. (*B*) Bar plot summarizing the Young’s modulus values showing that CENP-A nucleosomes are more elastic than H3 nucleosomes but become stiffer upon addition of CENP-C^CD^ (two-sided t-test *p*<0.0001). ∼1000 force curves were measured per condition.

**Table 1.** Nanomechanical force spectroscopy indicates that CENP-C^CD^ stiffens and suppresses CENP-A nucleosomal elasticity. Either H3 or CENP-A nucleosomes were *in vitro* reconstituted on plasmid DNA and imaged in fluid in the presence or absence of 2-fold or 4-fold excess CENP-C^CD^. Values were rounded up to 1 decimal point. N = number of nucleosomal particles measured. FC = number of force curves measured. For each condition, at least three independent replicates were performed (*SI Appendix*, Raw Data S1).

| Nucleosome | N | FC | Young's modulus (MPa) | Height (nm) | Diameter (nm) | Volume (nm<sup>3</sup>) |
| --- | --- | --- | --- | --- | --- | --- |
| Mononucleosomes |  |  |  |  |  |  |
| H3 | 5 | 24 | 35.4±13.9 | 5.2±0.53 | 11.3±1.2 | 371±107 |
| CENP-A | 4 | 34 | 18.5±15.6 | 5.7±0.53 | 11.7±2.3 | 387±86 |
| Nucleosome arrays |  |  |  |  |  |  |
| H3 | 48 | 997 | 11.3±4.1 | 3.8±0.3 | 14.0±1.2 | 393±68 |
| CENP-A | 46 | 977 | 5.8±3.0 | 3.7±0.3 | 13.7±1.0 | 370±61 |
| + 2X CENP-C<sup>CD</sup> | 48 | 1000 | 9.4±5.8 | 4.1±0.5 | 13.5±0.9 | 394±61 |
| + 4X CENP-C<sup>CD</sup> | 50 | 1014 | 15.2±10.5 | 4.1±0.6 | 14.0±1.2 | 426±61 |

Using these standardized nucleosomes, we then measured nucleosomal elasticity (*Methods*). First, consistent with the computational model, CENP-A and H3 mononucleosomes display uniform elasticity across their surfaces, behaving as homogenous cylinders (Fig. 2). Second, individual CENP-A nucleosomes are twice as elastic compared to H3 nucleosomes (18.5±15.6 MPa vs 35.4±13.9 MPa, respectively, Table 1).

*In vivo* nucleosomes exist in arrays. Therefore, we extended these experiments to arrays of nucleosomes reconstituted on 601-containing plasmids under identical conditions (*Methods*). As noted above, MNase digestion and AFM measurements confirmed that nucleosome arrays were reconstituted efficiently (*SI Appendix*, Fig. S3). Remarkably consistent with our computational results (Fig. 1*B*), and with the result for mononucleosomes (Table 1), the effective Young’s moduli of H3 and CENP-A nucleosomes are distinct. The Young’s modulus of H3 nucleosomes is 11.3±4.1 MPa, whereas CENP-A nucleosomes are nearly twice as elastic, at 5.8±3.0 MPa (Fig. 3*B*, Table 1).

### CENP-C^CD^ stiffens CENP-A nucleosomes *in vitro*

Our *in silico* experiments predicted that CENP-C^CM^ suppresses CENP-A nucleosomal motions and consequently innate elasticity (Fig. 1). We tested this prediction *in vitro*. We first examined the behavior of CENP-A nucleosomes in the presence of human or rat CENP-C^CM^. We observed a qualitative increase in cross-array clustering of CENP-A chromatin arrays (*SI Appendix*, Fig. S4). This rapid clustering by the CENP-C^CM^ fragment made it challenging to measure the rigidity of individual nucleosomes reliably. To resolve this challenge, we continued our investigation with CENP-C^CD^ which, as noted above, has the conserved binding motif of CENP-C^CM^ (*SI Appendix*, Fig. S2). The addition of human CENP-C^CD^ resulted in a 0.4 nm height increase of CENP-A nucleosomes (3.7±0.3 nm vs. 4.1±0.5 nm, Table 1, *SI Appendix*, Fig. S5, S6, Table S1), lending confidence that CENP-C^CD^ is bound to CENP-A nucleosomes.

Next, we measured the Young’s moduli of CENP-C bound vs. free CENP-A nucleosomes (*Methods*). With the addition CENP-C^CD^ at 2-fold excess, we observed that half the CENP-A nucleosomes remained highly elastic (∼5 MPa), but the other half lost elasticity by a factor of three (∼14.5 MPa) (Fig. 3*B*, Table 1, t-test *p*=0.015). One obvious interpretation of this distribution is that it arises from two distinct CENP-A sub-species: unbound and flexible versus bound and rigidified by CENP-C. To test this idea, we doubled the amount of CENP-C^CD^ to 4-fold excess. Under these conditions, virtually all CENP-A nucleosomes become stiffer (15.2±10.6 MPa, Table 1, *SI Appendix*, Fig. S6, S7).

These data show that *in silico,* and *in vitro* CENP-A nucleosomes possess innate elasticity and that CENP-C effectively suppresses the freedom of motions of CENP-A nucleosomes. From a thermodynamic perspective, elastic particles possess higher configurational entropy (33–35). In other words, elastic particles tend to be less ordered. Thus, we were curious to test whether nucleosomes with a broadened range of configurational states might collectively form less ordered chromatin and energetically disfavor compaction.

### CENP-C induces cross-array clustering *in vitro*, *ex vivo*, and *in vivo*

We first sought to tease out this idea by incubating *in vitro* reconstituted CENP-A chromatin arrays with or without CENP-C^CD^ and observed these arrays by in-air AFM. Upon addition of CENP-C^CD^, CENP-A arrays demonstrated a qualitative increase in cross-array clustering (Fig. 4*A*). This clustering was not observed for controls, namely CENP-C^CD^ incubated with either H3 chromatin or naked DNA (Fig. 4*A*).

**Figure 4.**
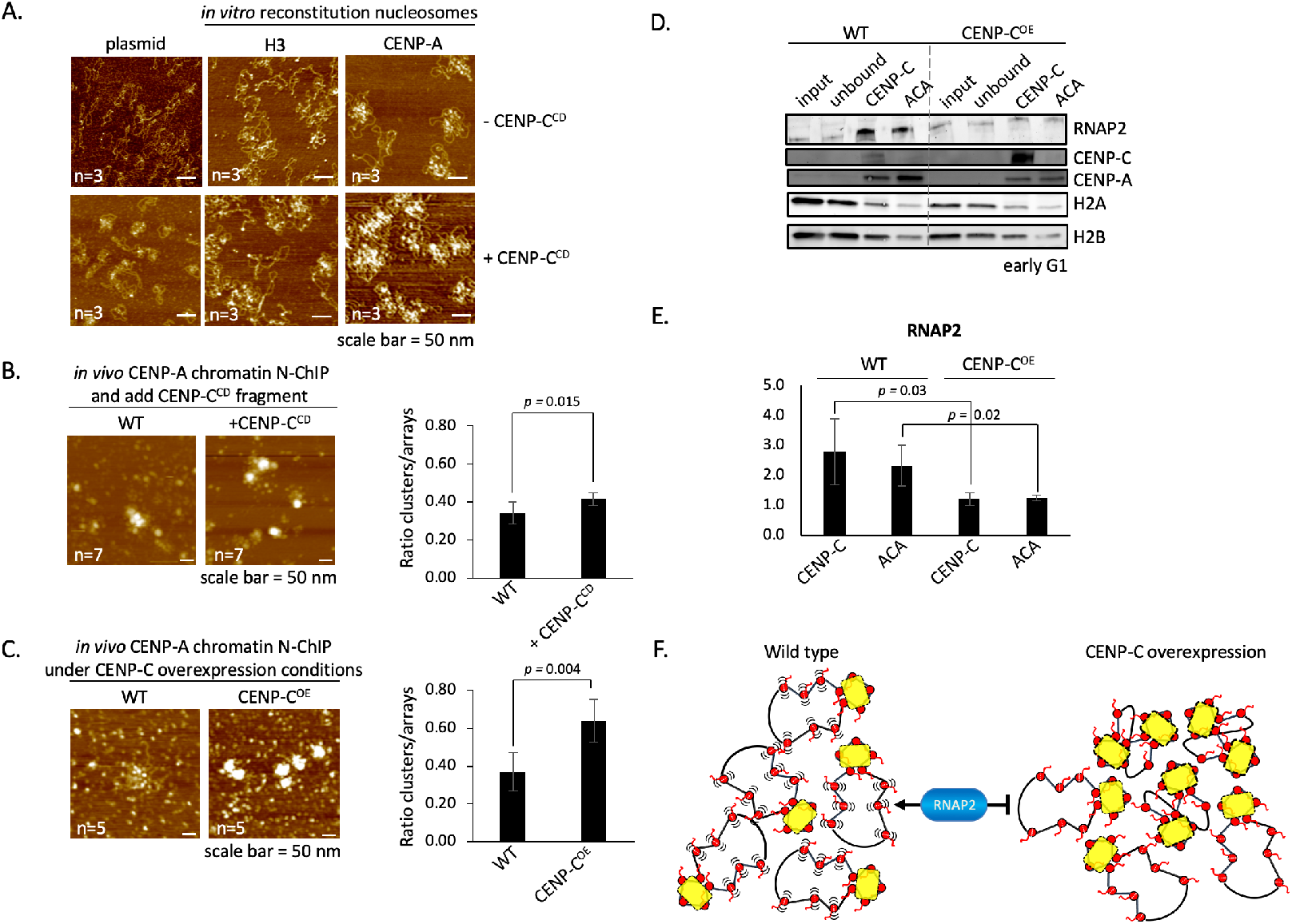
CENP-C overexpression compacts CENP-A chromatin, making it inaccessible to RNAP2. (*A*) Qualitative assessment of *in vitro* reconstituted chromatin showed that only CENP-A chromatin clustered in the presence of CENP-C^CD^ fragment. (*B*) To determine if the CENP-C^CD^ fragment used in the *in vitro* experiments could induce CENP-C chromatin compaction, we added CENP-C^CD^ for 30 minutes to isolated free CENP-A chromatin from HeLa cells. Compacted chromatin was scored over the total number of nucleosome arrays. (*C*) Similar analysis was performed on unbound CENP-A chromatin extracted from cells that either did (CENP-C^OE^) or did not (WT) overexpress CENP-C. (*D*) Western blot of serial N-ChIP probing for RNAP2 and various centromere and chromatin markers. Quantification of (*E*) RNAP2 levels were determined by LiCor’s software. The bar graphs represent three independent experiments. (*F*) Working model of CENP-C (yellow) overexpression inducing CENP-A chromatin (red) cross-array clustering thereby reducing access to RNAP2 (blue).

We next tested whether *ex-vivo*, kinetochore depleted CENP-A chromatin purified from human cells (*Methods*) (36) would cluster solely upon the addition of recombinant CENP-C^CD^ (Fig. 4*B*). Relative to free CENP-A chromatin, we observed a modest 1.2-fold increase in chromatin clusters upon CENP-C^CD^ incubation (34±6% vs. 42±4%, two-sided t-test *p*=0.015, Fig. 4*B*, *SI Appendix*, Table S2).

A logical hypothesis arising from these *in vitro* and *ex vivo* results is that excess CENP-C induces a more compact CENP-A chromatin state *in vivo*. To test this idea, we overexpressed full-length CENP-C (CENP-C^OE^) in human cells for three days, after which we purified kinetochore-depleted CENP-A chromatin by serial N-ChIP (*Methods*) (36). We quantified native CENP-A chromatin clusters using the same method as above. Upon CENP-C^OE^, we observed a nearly two-fold increase in chromatin clusters relative to the wild-type control (37±10% vs. 64±11%, two-sided t-test 0.004, Fig. 4*C*, *SI Appendix*, Table S2). Thus, *in vitro*, *ex vivo*, and *in vivo*, CENP-C increases the population of CENP-A chromatin clusters.

### CENP-C overexpression limits centromeric chromatin accessibility *in vivo*

It has been demonstrated that chromatin accessibility is prognostic of transcriptional competency across the genome (37, 38). This correlation was first reported decades ago in two seminal works demonstrating nuclease hypersensitivity of actively transcribing loci (39, 40). We hypothesized that an innately open CENP-A chromatin state would be accessible, whereas excess CENP-C should reduce the accessibility of CENP-A chromatin *in vivo*. One read-out of altered compaction status would be reduced accessibility of CENP-A chromatin to transcriptional machinery.

To test this idea, we performed CENP-C^OE^ for three days and synchronized the cells to early G1, when centromeres are transcribed in human cells (13, 14, 41). From these cells, we purified CENP-C bound centromeric chromatin as well as any residual CENP-A chromatin by serial N-ChIP (*Methods*) (36). We assessed the occupancy of active RNA polymerase 2 (RNAP2) on these purified native chromatin arrays from wild type or CENP-C^OE^ cells. By western blot analysis, when CENP-C is over-expressed, we observed a significant reduction in RNAP2 levels on centromeric chromatin (3- and 2-fold reduction, resp.; two-sided t-test *p*<0.05; Fig. 4*D*, *E*, *SI Appendix*, Table S3). Thus, CENP-C overexpression leads to both, CENP-A chromatin clustering, and reduced accessibility of transcriptional machinery.

These data show that CENP-C overexpression suppresses accessibility of centromeric chromatin (Fig. 4F), correlating with the attenuation of transcriptional machinery.

## Discussion

Not all nucleosomes are identical, as many contain histone variants, giving them distinct structures and functions (1, 2, 6). In this report, we systematically teased apart how a single histone variant encodes mechanical properties to its nucleosome, which were dramatically modified by a small fragment of its cognate protein partner. Using novel *in silico* computational modeling and *in vitro* single-molecule nanoindentation force spectroscopy, we directly measured effective elasticity of nucleosomes and found that CENP-A is more elastic than canonical nucleosomes (Fig. 1-3, Table 1). Indeed, we found remarkable agreement between the computation model to derive the Young’s modulus, and the experimental data measuring the elasticity. Second, our findings of noticeably elastic CENP-A nucleosomes have important implications. Specifically, one expects from general statistical physics reasoning that CENP-A nucleosomes contain excess entropy compared to canonical nucleosomes, which, in turn, will be lost upon formation of compacted chromatin. Hence, one may anticipate extra entropic resistance to compaction for chromatin enriched with CENP-A nucleosomes.

CENP-C is the essential CENP-A binding protein, which facilitates the assembly of the kinetochore (42–44), and has been shown to alter local CENP-A nucleosomes dynamics (16–18). Previous FRET and hydrogen/deuterium exchange mass spectrometry experiments focused on how CENP-C^CD^ binding alters internal CENP-A mononucleosome dynamics. These data show that human CENP-C^CD^ restricts DNA gyre gapping, sliding, and protects the internal H4/H2A interface (16–18). In our prior computational modeling we showed that CENP-A samples broadened conformational states (15). From this, we predicted that CENP-C limits configurations of CENP-A. Indeed, when we modeled CENP-A nucleosomes alone, vs. those bound to CENP-C^CM^, we observed marked diminution of nucleosome motions, and increased Young’s moduli, representing lost conformational flexibility (Fig. 1*B*, *SI Appendix*, Fig. S1). Direct elasticity measurements by nanoindentation force spectroscopy revealed that CENP-C^CD^ stiffens the CENP-A nucleosomes in a dose-dependent manner. A physical analogy is that CENP-C behaves as a nanoscale staple on the surface of the CENP-A nucleosome, inhibiting intra and intermolecular motions and propagates to the chromatin fiber. Furthermore, the homodimerization domain of CENP-C likely exaggerates cross-array clustering (45). Thus, a speculative prediction from this model is that the centromeric fiber harbors a free CENP-A domain to allow cell-cycle regulated transcription of centromeres—required for the loading of new centromeric proteins (13, 14, 41).

We note that CENP-C expression is tightly regulated, despite over-expression of many centromere proteins in human cancers, including CENP-A (31, 46–48). Taken in the context of our findings in this report, maintaining the correct ratio between CENP-A and CENP-C *in vivo* might be critical for preserving the mechanical features of centromeric chromatin. In human cancer cells, where CENP-A is over-expressed and ectopically localized to subtelomeric breakpoints (31, 46), one unexpected mechanical outcome might be the induction of large swathes of elastic CENP-A chromatin at inappropriate regions of the genome (49). This will be an exciting avenue to pursue in future studies.

Centromeric DNA and centromeric protein genes are rapidly evolving (12, 50–55). Not all species share all kinetochore components: centromeric genes are lost, duplicated, and sometimes invented (56–58). Despite these evolutionary changes, the distinctive chromatin structure of centromeres must be maintained. Investigating whether CENP-A elasticity is conserved or co-evolve with kinetochore proteins, will shed light on centromeric evolution. The centromere will then in turn serve as an excellent model to study the evolution of epigenetic systems.

## Methods

### All-atom computational modeling

We built three nucleosomal systems for simulation: the CENP-A nucleosome as described previously (59). and the CENP-A nucleosome with one and two CENP-C central domain fragment bound from PDB ID: 4X23(19). The CENP-C^CD^ fragments were docked onto the CENP-A interface using the CE algorithm(60) of PyMOL (The PyMol Molecular Graphics System). We set up both systems to initiate from the final time point of our previous 2 μs simulation and the coordinates, velocities, parameters, and system setup and analysis methods were replicated (59). For quality control and checks for equilibration, the energy minimization, equilibration, and running RMSD for the simulations (*SI Appendix*, Fig. S8). Both CENP-A and CENP-A with one and two CENP-C^CD^ bound (19) ran for an additional microsecond and the first 600 ns of simulation time were truncated from the dataset for further analysis and to account for equilibration. For a control to compare to this dataset, we also analyzed the H3 nucleosome from our previous work (15). In addition to our prior description, after energy minimization we checked our structures for potential clashes based on van der Waals radii through the accepted range of 0.4 – 1.0 Å and verified that there were no clashes in the nucleosome structures.

Furthermore, we calculated the relative positions of three phosphate backbone atoms at positions −33, −43, and +38 numbered from the 5’ (−) to 3’ (+) direction relative to the pseudo-dyad. The distances between these points and the skew of the triangle formed were measured and then plotted with the initial position of residue −33 set to (0, 0) on a xy-plane. The distribution of Δy and Δx of +38 relative to −33 and −34 was used to measure DNA gaping and sliding respectively. We visualized these distributions with standard box plots showing the mean, the interquartile range, and whiskers extending to the extrema. The distribution of polygons contains the minima and maxima of all three vertices were plotted visually with triangles to present changes in skew and the range of sizes. We executed RMSF and center-of-mass motion calculations as described previously (59).

### *In silico* calculation of Young’s Modulus

The goal of this analysis is to model each nucleosome as a homogenous elastic “minimal” cylinder for each time step of the simulation, retrieve the cylinder height and radius distributions, and from this data calculate the in silico Young’s Modulus of the nucleosomes. Our method to calculate the dimensions of the minimal cylinders follows the workflow:

[1] Orient the nucleosomes so that they lie “flat” on the x-y plane. To achieve this, we calculated the principal axes of the moment of inertia, where the first principal axis defines the broadest plane of the nucleosome. The axes of symmetry of the nucleosomes align with the three principal axes, p_1_, p_2_, p_3_, with the center-of-mass at the origin.

[2] Calculate the surfaces of the cylinder so that they coincide with stiffer regions of the nucleosomes. We addressed this issue by calculating the root mean square fluctuations (RMSF) of each residue along the simulation since the structural disorder of a region positively correlates with local structural fluctuations. Since RMSF is a time-averaged parameter, multiple timesteps are required to calculate fluctuations of residues. As a result, we divided the simulation into windows (800 windows per simulation) and calculated the RMSF for each residue in each window.

[3] Retrieve the average heights, radii, and the variances of these distributions. To do so, we sorted the C-α coordinates by their z-axis coordinates and selected the z coordinate of the residue where ten stiffer residues below an RMSF threshold were excluded outside of the cylinder volume. From the height, h, and radius, r, data we calculated the average h and r, the variance or spread of the distributions, and the standard deviations Δr and Δh.

[4] The outputs from step [3] then served as the variable inputs to calculate the Young’s Modulus of each system. The work done in the deformation of an elastic material is stored in the form of strain energy which we calculate for the deformation of the cylinder in the absence of the shear stresses. In our simulations, the amplitude of vibrations depends on the amount of energy given to the system from the temperature, or the thermal bath of the solvent. From equipartition theorem, 1/2 k_b_T (where k_b_ is the Boltzmann constant and T is temperature, 300 K) is the amount of energy attributed to the observed cylinder deformation. From the data on the average cylinder conformation, the magnitude of elastic deformation, and the energy input from the thermal bath we calculate the Young’s modulus.

### Single-molecule nanoindentation force spectroscopy of mononucleosomes

H3 (H3 mononucleosome on 187bp of 601 sequence cat#16-2004, EpiCypher, Research Triangle Park, NC) and CENP-A mononucleosome (CENP-A/H4 cat#16-010, H2A/H2B cat#15-0311, 187bp of 601-sequence cat#18-2003, EpiCypher, Research Triangle Park, NC) samples were diluted 1:5 in 2 mM NaCl with 4 mM MgCl (pH7.5) and deposited onto freshly cleaved mica that had previously been treated with aminopropyl-silantrane (APS) as described (32, 61, 62). Samples were incubated on mica for ∼3 minutes, excess buffer was rinsed with 400 μL ultrapure, deionized water, and gently dried under an argon stream. Imaging was performed with a commercial AFM (MultiMode-8 AFM, Bruker, Billerica, MA) using silicon-nitride, oxide-sharpened probes (MSNL-E with nominal stiffness of 0.1 nN/nm, Bruker, Billerica, MA). Deposited sample was rehydrated with 10 mM HEPES (pH 7.5), 4 mM MgCl. Imaging was performed in AFM mode termed “Peak-Force, Quantitative NanoMechanics” or PF-QNM. Images were preprocessed using the instrument image analysis software (Nanoscope v8.15) and gray-scale images were exported to ImageJ analysis software (v1.52i). First nucleosomes were identified as described (32, 61) and subsequently roundness was determined. The Young’s modulus was determined by the instrument image analysis software (Nanoscope v8.15).

### Optimization of single-molecule nanoindentation force spectroscopy

Nucleosomes that lay flat, have a round appearance, whereas nucleosomes laying on their side would have an oval appearance. We measured the roundness of both H3 and CENP-A mononucleosomes and found that almost all nucleosomes had a round appearance (*SI Appendix*, Fig. S2*A*).

The use of AFM nanoindentation of nucleosomes raise two more concerns. One is that the size of the probe is of the same order of magnitude as the nucleosome. Therefore, widely-used, Hertz-type models used to extract elasticity from indentation data would only provide an effective elasticity that depends on the indentation geometrical parameters such as probe size and precise point of indentation on the nucleosome. This effective elasticity would, however, be comparable between the two types of nucleosomes and their relative values would be comparable to those obtained *in-silico.* The probe sizes used did not vary significantly but we needed to address the possibility that the extracted elasticity depends strongly on the exact point of indentation. If the nucleosome is not uniformly elastic, the precise position of the AFM probe tip could be a critical factor. If the nucleosomes are uniformly elastic, slight differences in where on the nucleosome the elasticity is measured would not be a major concern. We therefore measured the Young’s modulus across mononucleosomes (*SI Appendix*, Fig. S2*B*). We found that, in general, effective elasticity did not vary significantly across nucleosomes (*SI Appendix*, Fig. S2*B*).

### Single-molecule nanoindentation force spectroscopy of nucleosome arrays

*In vitro* reconstitution of CENP-A nucleosome arrays (CENP-A/H4 cat#16-010 and H2A/H2B cat#15-0311, EpiCypher, Research Triangle Park, NC) and H3 (H3/H4 cat#16-0008 and H2A/H2B cat#15-0311, EpiCypher Research Triangle Park, NC) on a 3kb plasmid containing a single 601 sequence (pGEM3Z-601 from Addgene #26656) were performed as previously described (32, 61). Human CENP-C_482-527_ fragment (CENP-C^CD^) (19) and rat CENP-C_710-740_ (CENP-C^CM^) (ABI Scientific, Sterling, VA) was added in 2.2-fold or 4-fold molar excess to CENP-A nucleosomes. Imaging was performed by using standard AFM equipment (Oxford Instruments, Asylum Research’s Cypher S AFM, Santa Barbara, CA). To be able to measure the Young’s modulus, the reconstituted chromatin was kept in solution containing 67.5 mM NaCl and 2 mM Mg^2+^ and Olympus micro cantilevers (cat# BL-AC40TS-C2) were used. Before each experiment, the spring constant of each cantilever was calibrated using both GetReal™ Automated Probe Calibration of Cypher S and the thermal noise method (63). Obtained values were in the order of 0.1 N/m. As a reference to obtain the indentation values, the photodiode sensitivity was calibrated by obtaining a force curve of a freshly cleaved mica surface. All experiments were conducted at room temperature. Force-curves for ∼50 nucleosomes for all three conditions were measured using both ‘Pick a Point’ and force-mapping mode. The maximum indentation depth was limited to ∼1.5 nm and the maximum applied force was 150-200 pN. For our analyses, we used Hertz model with spherical indenter geometry for Young’s Modulus measurements, δ = [3(1 – ν^2^)/(4ER^1/2^)]^2/3^F^2/3^ (for a spherical indenter), where ν is the Poisson ratio of the sample, which is assumed to be 1/3 as in studies reported previously (24, 27); δ, F, E, and R are the indentation, force, Young’s modulus of the sample and radius of the tip respectively. The radius of the tip was confirmed by SEM and found to be about 10 nm in width. Graphs were prepared using ggplot2 package for R.

### AFM and cluster analysis

Imaging of CENP-C and CENP-A N-ChIP and bulk chromatin was performed as described (32, 61)with the following modifications. Imaging was acquired by using commercial AFM equipment (Oxford Instruments, Asylum Research’s Cypher S AFM, Santa Barbara, CA) with silicon cantilevers (OTESPA or OTESPA-R3 from Olympus with nominal resonances of ∼300 kHz, stiffness of ∼42 N/m) in noncontact tapping mode or commercial AFM (MultiMode-8 AFM, Bruker, Billerica, MA) using silicon cantilevers (OTESPA or OTESPA-R3 from Olympus). Reconstituted H3 or CENP-A chromatin with or without rat or human CENP-C^CM^ or human CENP-C^CD^ fragments or on kinetochore-depleted chromatin obtained from HeLa cells (36) with and without human CENP-C^CD^ was deposited on APS treated mica In addition, kinetochore-depleted chromatin obtained from HeLa cells (36). APS-mica was prepared as previously described (32, 61). The samples were incubated for 10 min, gently rinsed with two times 200 μL ultra-pure water, and dried with inert argon gas before imaging.

For the compaction study, we added 1 ng CENP-C fragment to purified ACA samples and incubated them for 30 minutes prior to deposition on APS-mica and subsequent imaging. To quantify the chromatin compaction, we manually counted chromatin clusters based on their size being at least twice as wide as an individual nucleosome, but with an identifiable entry and exit DNA strand, over the total number of nucleosome arrays.

### Native Chromatin-Immunoprecipitation and western blotting

Human cell line HeLa were grown in DMEM (Invitrogen/ThermoFisher Cat #11965) supplemented with 10% FBS and 1X penicillin and streptomycin cocktail. N-ChIP experiments were performed without fixation. After cells were grown to ∼80% confluency, they were harvested as described here (27, 36, 59), but with a few modifications. These were that all centrifugation was done at 800 or 1000 rpm at 4°C. Chromatin was digested for 6 minutes with 0.25 U/mL MNase (Sigma-Aldrich cat #N3755-500UN) and supplemented with 1.5 mM CaCl_2_. The first N-ChIP was with 5 μL guinea pig CENP-C antibody, subsequently the unbound fraction was N-ChIP’ed with 5 μL ACA serum (*SI Appendix Methods*). For CENP-C overexpression we transfected HeLa cells with pEGFP-CENP-C using the Amaxa Cell Line Nucleofector Kit R (Lonza cat#VVCA-1001) per manufacturer’s instructions. HeLa cells were synchronized to early G1 by double thymidine block (0.5 mM, Sigma-Aldrich cat#T9250). After the first block of 22 hours, cells were released for 12 hours, followed by a second thymidine block of 12 hours. Cells were released for approximately 11 hours, which corresponds to early G1, based on our previous reports (27, 59).

### Quantification and Statistical Analyses

Significant differences for nucleosome height measurement from AFM analyses and significant differences for immunostaining quantification, and chromatin compaction quantification, were performed using the 2-sided t-test as described in the figure legends and main text. Significant differences for the Young’s modulus of *in vitro* reconstituted H3, CENP-A, and CENP-A + CENP-C^CD^ were determined using 1-way ANOVA test using GraphPad Prism software. Significance was determined at *p*<0.05.

## Supporting information

Raw Data S1

## Authors contribution

Conceptualization: DPM, MP, TR, GAP, and YD; Methodology: DPM, TR, MP, MB, EKD, GAP, and YD; Investigation: DPM, TR, MP, MB, and EKD; Writing: DPM, MP, GAP, and YD.; Funding Acquisition: EKD, GAP, and YD; Visualization: DPM, MP, and EKD; Supervision: GAP and YD.

## Acknowledgements

We thank Tom Misteli, Sam John, and members of our laboratory for critical comments; Will Heinz for discussions on force spectroscopy experiments; Carlos S Floyd for independent validation of *in silico* Young’s modulus calculations; Stephan Diekmann for gifting GFP-CENP-C; and Yawen Bai for advice on CENP-C fragments and binding assays and gifting human CENP-C^CM^. YD thanks participants of the GRC Chromosome Dynamics for very useful and positive feedback on this work. DPM, TR, MP, BM, EKD, and YD are supported by the Intramural Research Program of the National Institutes of Health. MP is supported by the joint NCI-UMD Cancer Technology Partnership. GAP is supported by NSF grant CHE-1800418.

## SI Appendix, Methods

### Key Resources Table

| **Antibody** | **Source** | **Identifier** | **Application** | **Quantity** |
| --- | --- | --- | --- | --- |
| ACA serum | BBI Solutions | SG140-2 | N-ChIP | 5 μL |
| α-CENP-A (mouse) | Abcam | ab13939 | IF | 1:1000 |
| α-CENP-A (rabbit) | Milipore | 04-205 | WB | 1:3000 |
| α-CENP-C (guinea pig) | MBL International | PD030 | N-ChIP | 5 μL |
| α-CENP-C (rabbit) | Santa Cruz | sc-22789 | IF, WB | 1:1000; 1:500 |
| α-RNA polymerase II | Abcam | ab5095 | WB | 1:500 |
| α-H2A | Abcam | ab18255 | WB | 1:1000 |
| α-H2B | Abcam | ab1790 | WB | 1:1000 |

IF = immunofluorescence; N-ChIP: native chromatin immunoprecipitation, WB = western blot

| **Software and Algorithms** |  |
| --- | --- |
| Gwyddion | http://gwyddion.net/ |
| NanoScope | http://www.nanophys.kth.se/nanophys/facilities/nfl/afm/icon/bruker-help/Content/SoftwareGuide/NanoScope815CoverPage.htm |
| Asylum Research | Version 15 |
| Igor Pro | https://www.wavemetrics.com/taxonomy/term/87 |
| R | https://www.r-project.org/ |
| ggplot2 | https://ggplot2.tidyverse.org/ |
| LiCor Image Studio | https://www.licor.com/bio/products/software/image_studio_lite/ |
| NIH ImageJ | https://imagej.nih.gov/ij/ |
| GraphPad Prism 8 | https://www.graphpad.com/ |
| Bio-Formats | https://www.openmicroscopy.org/bio-formats/ |
| PyMOL | https://pymol.org/2/ |
| MEME | http://meme-suite.org/tools/meme |

### Contact for Reagent and Resource Sharing

Requests for further information or reagents should be directed to the Lead Contacts: Yamini Dalal and Garyk Papoian.

---

**Figure S1.**
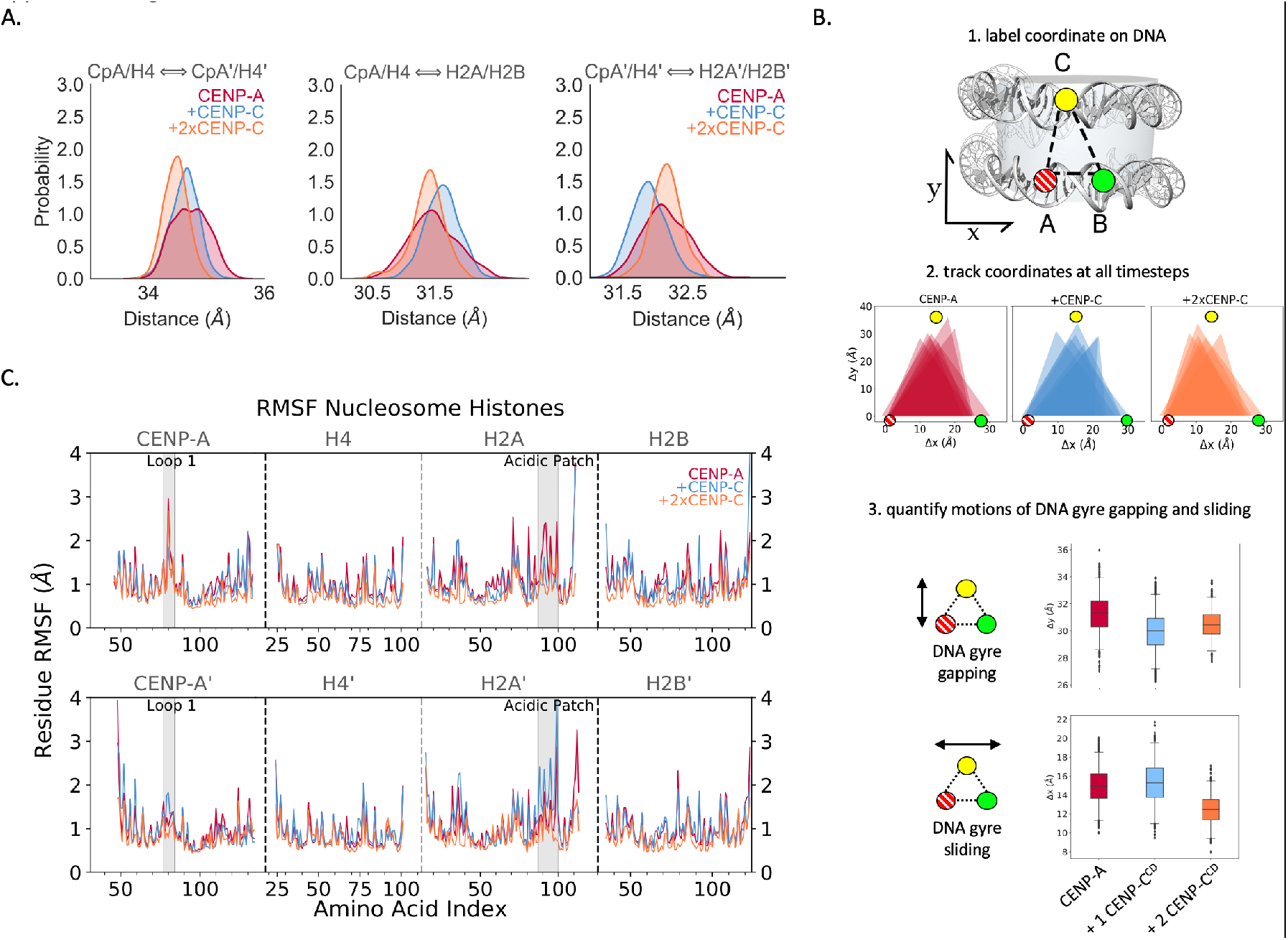
Two CENP-C^CD^ fragment strengthens stifening of CENP-A nucleosomes. (*A*) The distance between the center of mass (COM) of histone dimers is shown in red for CENP-A, blue for CENP-A + 1 CENP-C^CD^, and in orang for CENP-A + 2 CENP-C^CD^. Two CENP-C^CD^ fragment exaggerated the COM distances compared to a single CENP-C^CD^ fragment, which means that 2 CENP-C^CD^ further induces a global loss of CENP-A nucleosome flexibility. (*B*) The free energy landscape of CENP-A nucleosomes alone or CENP-A nucleosome with CENP-C^CD^ fragment was determined by principle component analysis. CENP-A nucleosomes display a rugged free energy landscape, which is locked down when CENP-C^CD^ is bound, increasing the connectivity of the energetic minima. All-atom computational modeling of DNA gyre gapping or DNA gyre sliding of CENP-A nucleosome alone or bound to either 1 or 2 CENP-C^CD^ fragments. (*C*) Residue root mean square fluctuations (RMSF) shows freezing of local flexibility in the CENP-A nucleosome shown in red, 1 CENP-C^CD^ bound shown in blue, and 2 CENP-C^CD^ bound shown in orange. In the region of CENP-C^CD^ binding, the first heterotypic half on the top panel, CENP-C is seen to freeze the acidic patch and the loop 1 region of CENP-A. One CENP-C^CD^ creates asymmetry, especially at the C-terminal end of H2A and H2B, this is abrogated when the second CENP-C^CD^ is bound. Dashed lines separate individual histones.

**Figure S2.**
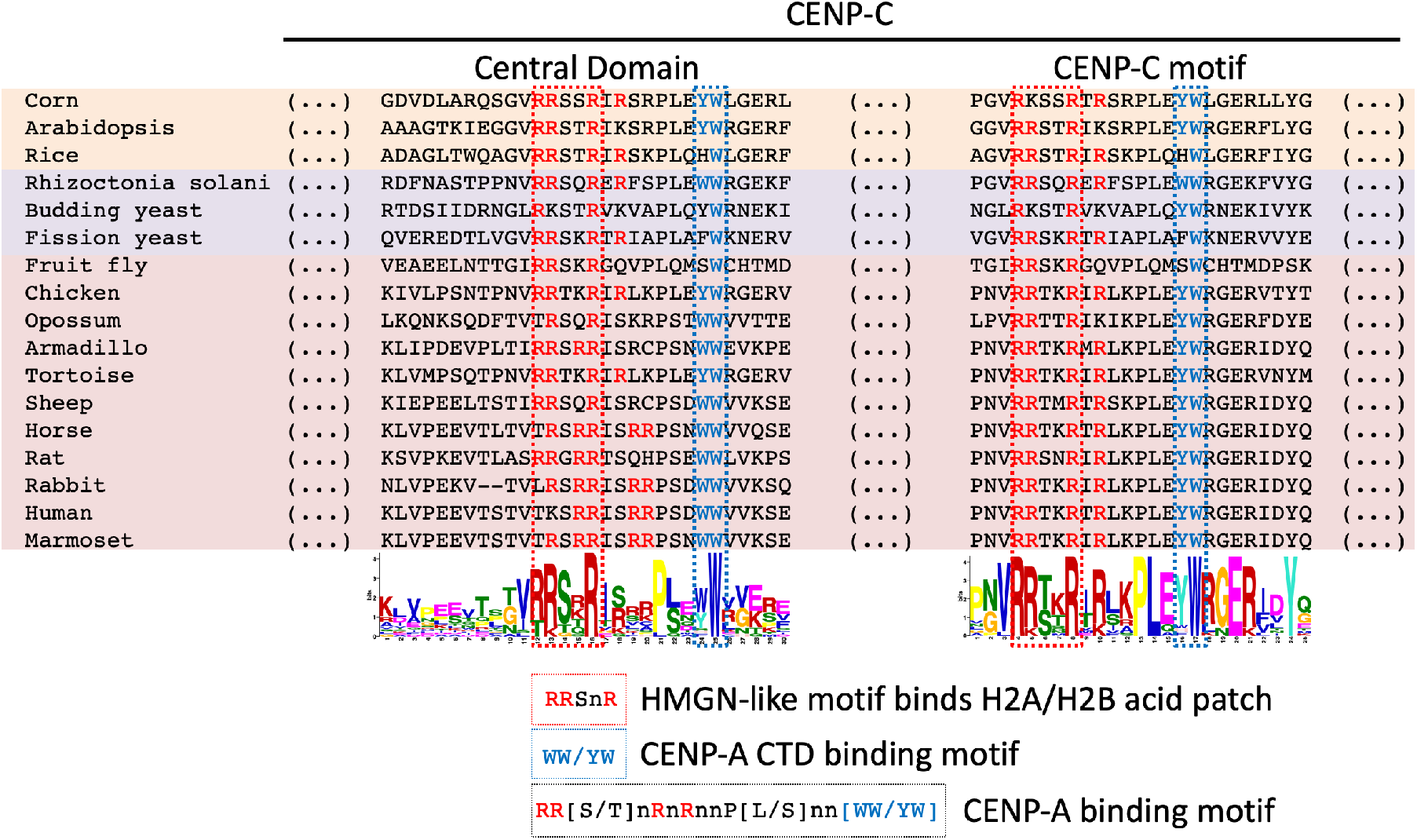
CENP-C^CM^ and CENP-C^CD^ have conserved CENP-A nucleosome binding motifs. Alignment of various plant, fungal, and animal species shows that within the poorly conserved central domain and the well conserved CENP-C motif the RR(S/T)nR motif and the WW/YW motif are highly conserved. These two motifs are separated by 7 residues, creating a conserved H2A/H2B acid patch and C-terminal tail of CENP-A binding motif (RR(S/T)nRnRnnP(L/S)nn(WW/YW).

**Figure S3.**
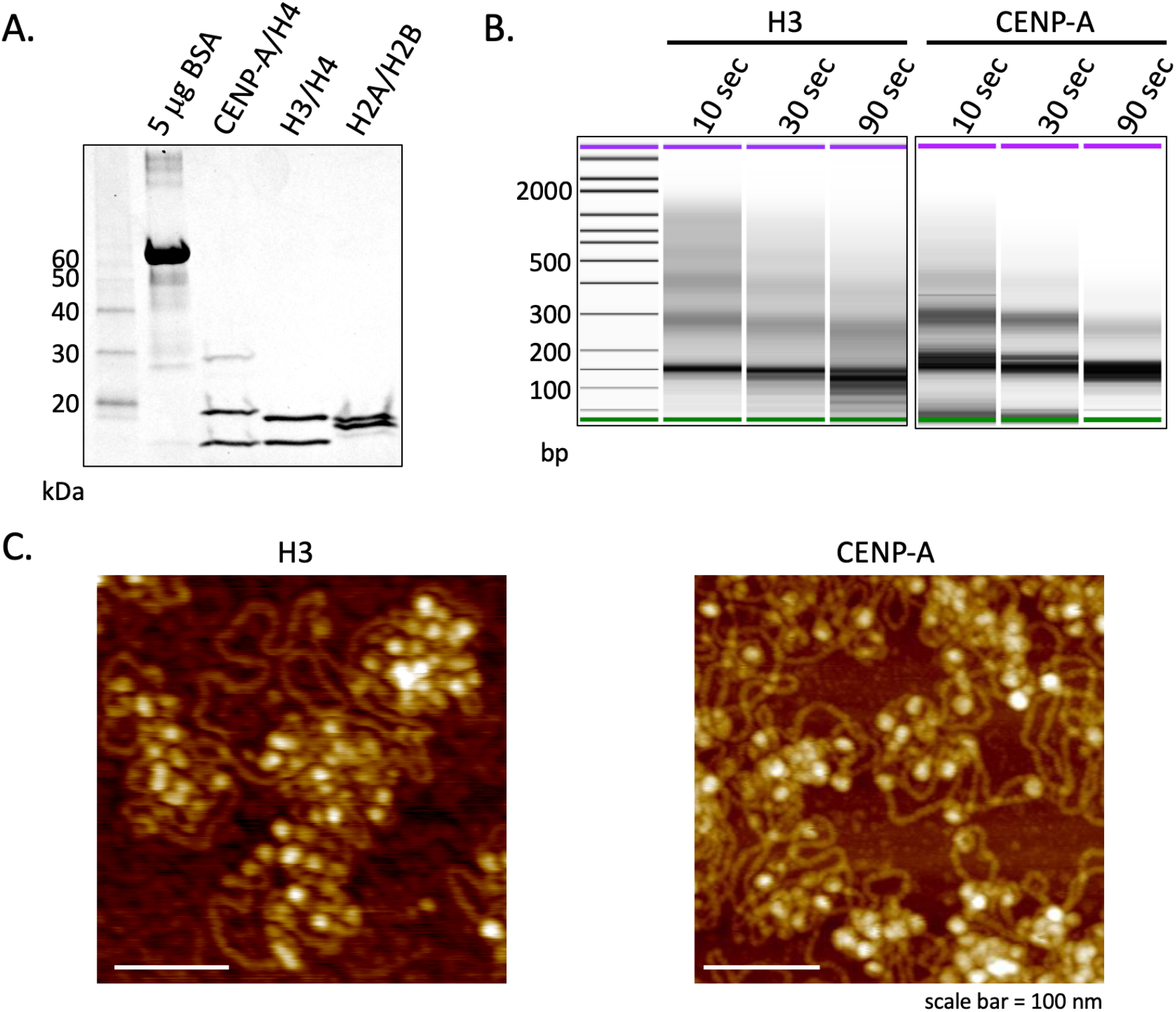
Nucleosomal arrays show MNase ladder. (*A*) Histone protein concentration was determined by Coomassie staining. (*B*) BioAnalyzer results from 10, 30, and 90 seconds of MNase digested reconstituted H3 and CENP-A nucleosomes on 3 kbp plasmids a classic chromatin ladder. (*C*) Representative in air AFM images of H3 and CENP-A reconstituted chromatin.

**Figure S4.**
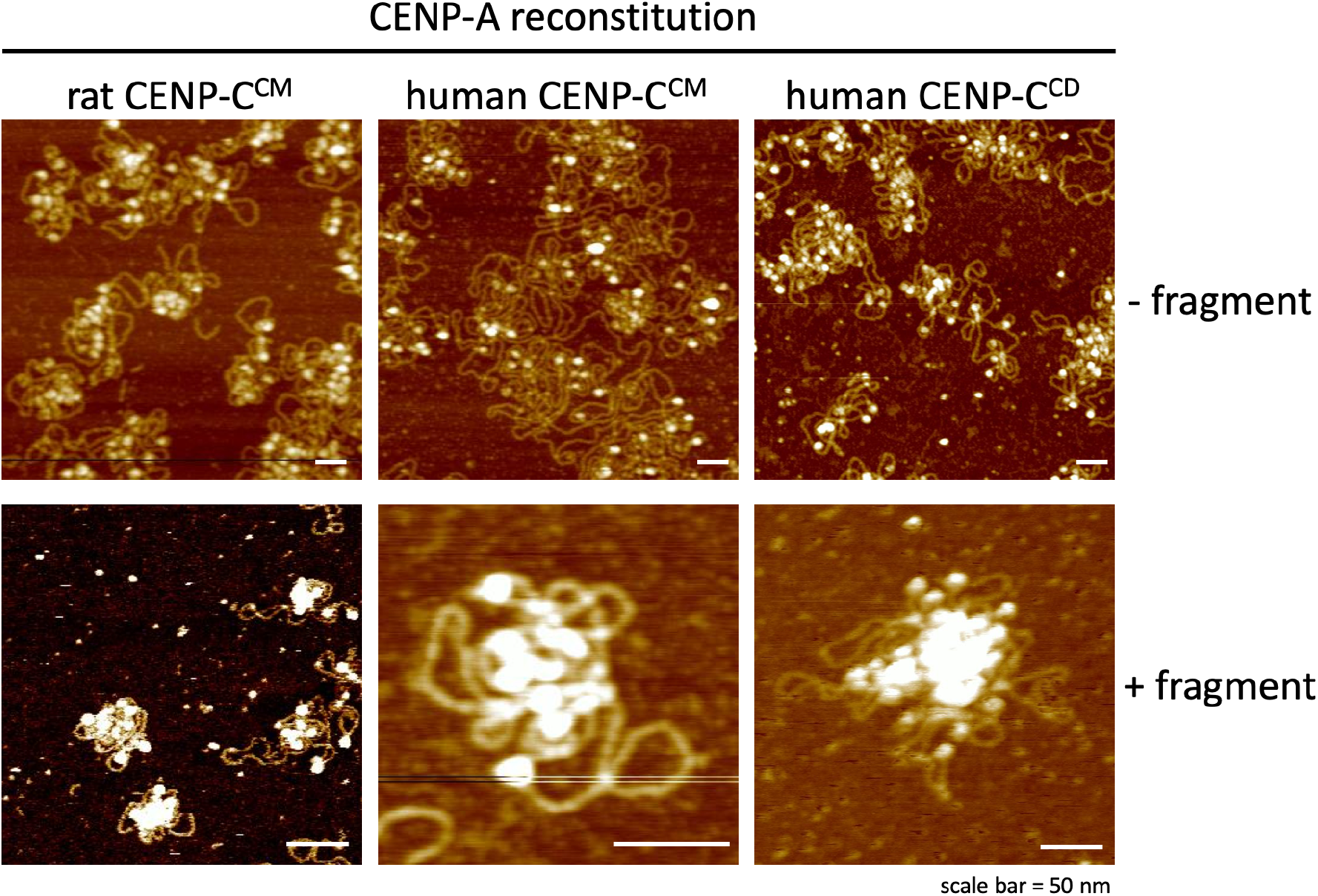
CENP-C induces CENP-A chromatin clustering. Reconstituted CENP-A chromatin was incubated with either rat CENP-C^CM^, human CENP-C^CM^, or human CENP-C^CD^ fragments for 30 minutes. Cluster formation was observed with all three fragments.

**Figure S5.**
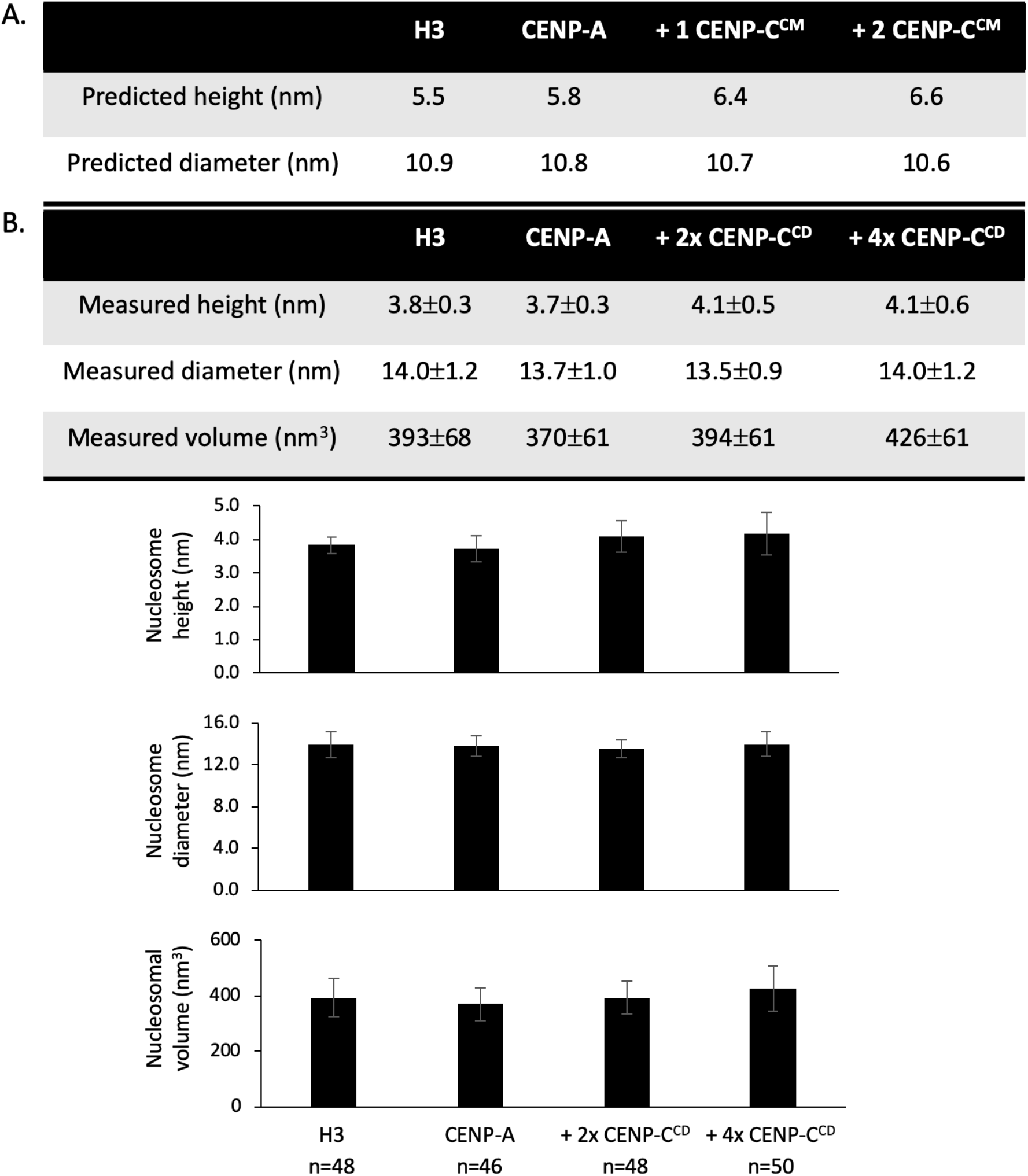
CENP-C^CD^ modestly increases CENP-A nucleosome heights. (*A*) Height and diameter predictions from the computational modeling experiment described in Figure 1A. (*B*) CENP-C^CD^ modestly increases height of *in vitro* reconstituted CENP-A nucleosomes H3 and CENP-A nucleosome were in vitro reconstituted, and by in fluid AFM, we measured their dimensions (height, diameter, and volume). The height distribution is shown in the violin plot containing a bar plot. CENP-A nucleosomes are ever so slightly smaller than H3 nucleosomes. The addition of CENP-C^CD^ fragment, which can only bind CENP-A nucleosomes, we observed an increase in height and in a dose-dependent manner its volume.

**Figure S6.**
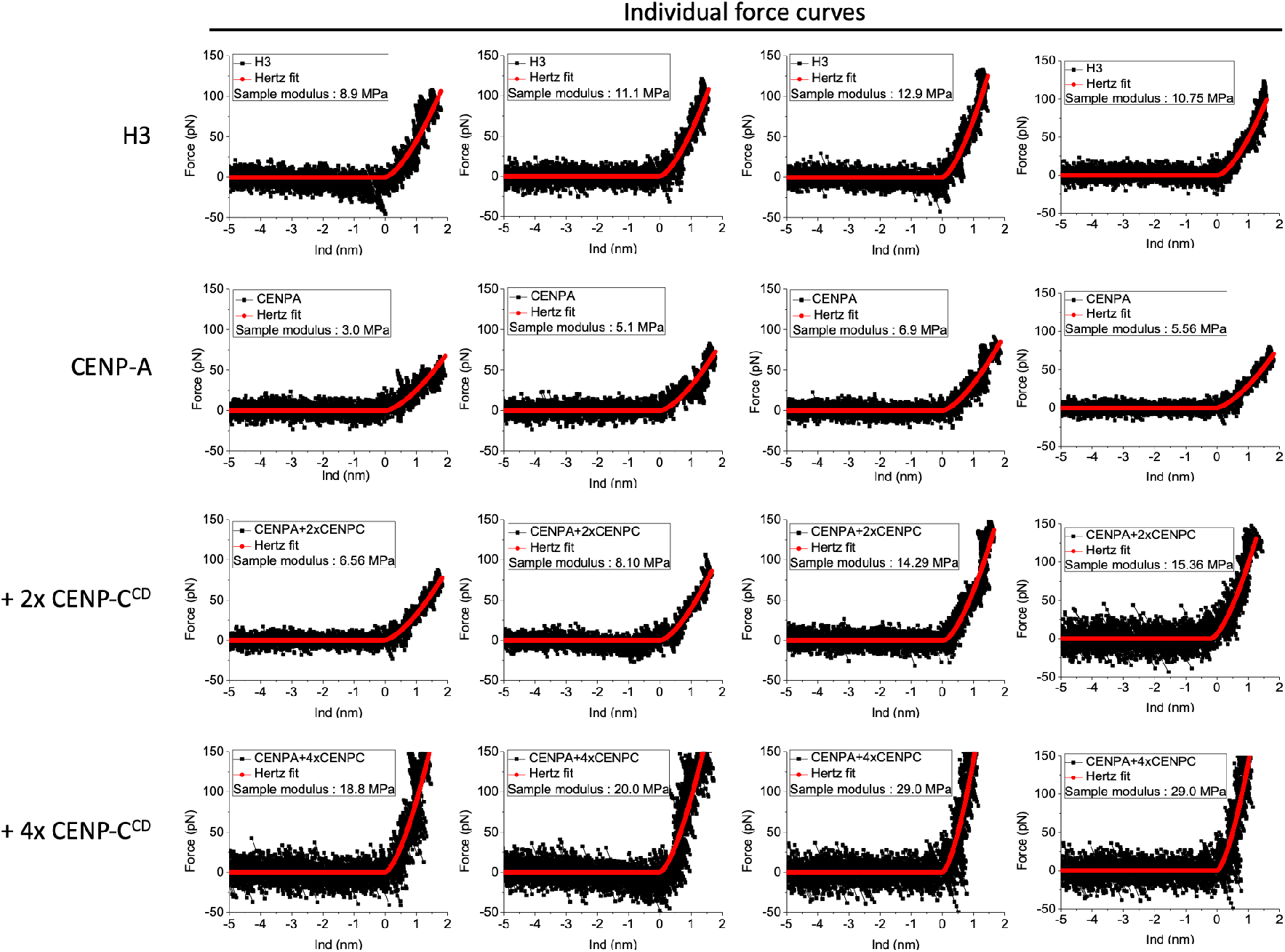
Examples of force curve measurements. Four representative force curves for H3 nucleosomes, CENP-A nucleosomes, CENP-A nucleosomes with 2-fold excess CENP-C^CD^ fragments, and CENP-A nucleosomes with 4-fold excess CENP-C^CD^ fragments are shown.

**Figure S7.**
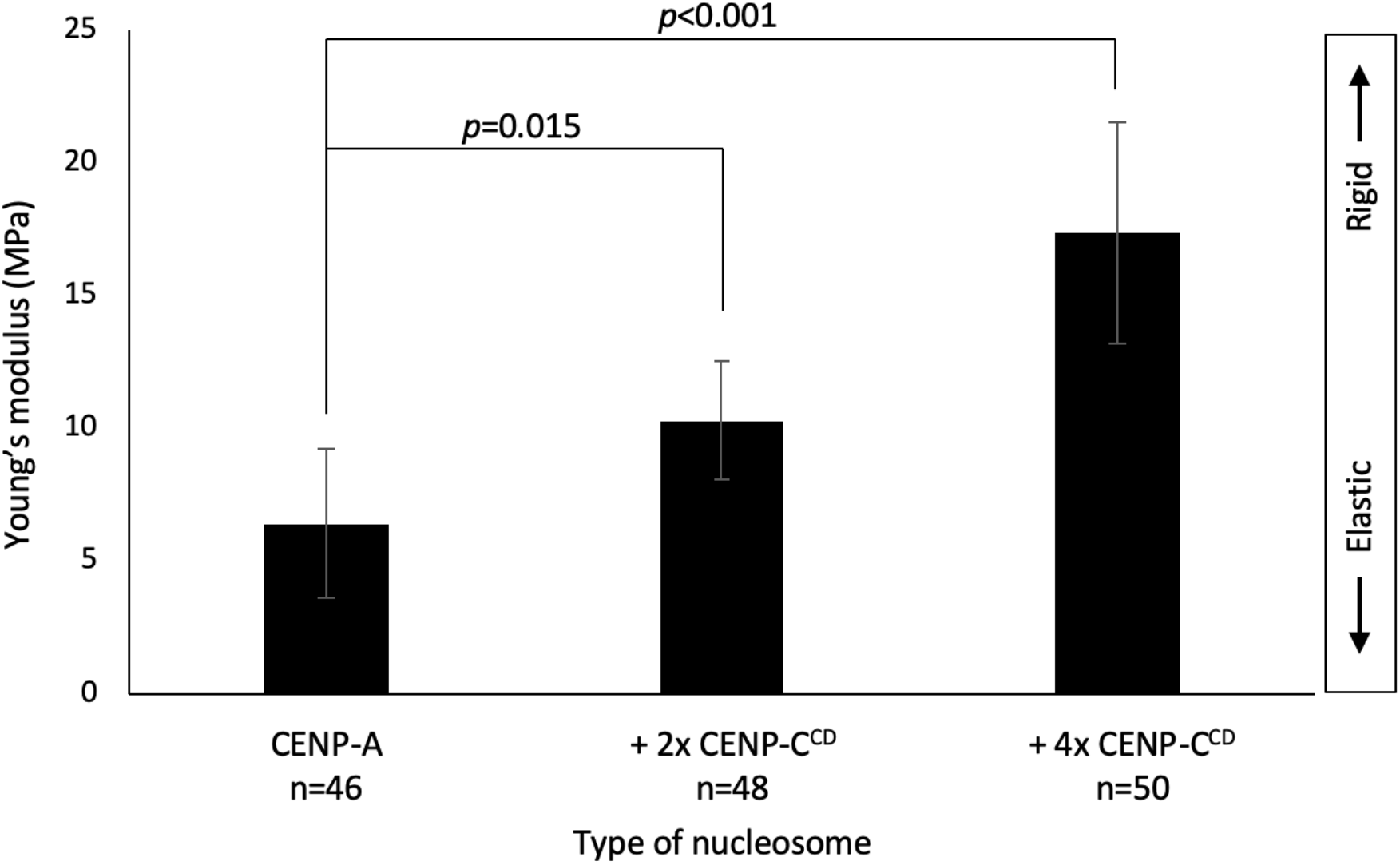
4-fold excess of CENP-C^CD^ results further increased CENP-A nucleosomes rigidification. Bar plot summarizing the Young’s modulus values showing that CENP-A nucleosomes become stiffer upon addition of 2-fold excess CENP-C^CD^ (two-sided t-test *p*=0.015), and even stiffer upon addition of 4-fold excess CENP-C^CD^ binding (two-sided t-test *p*<0.001). ∼1000 force curves were measured per condition.

**Figure S8.**
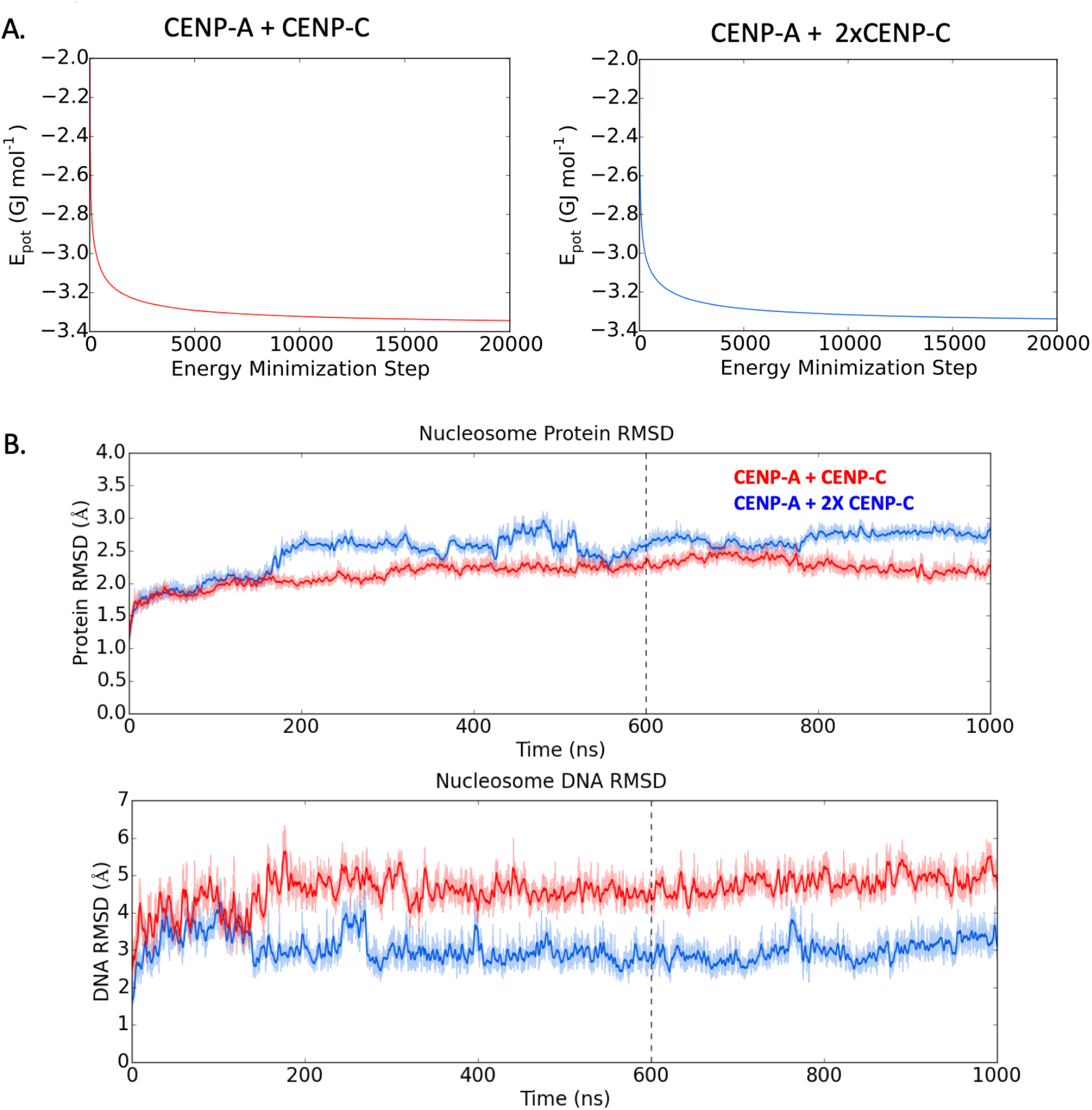
Quality control of computational simulations. (*A*) The systems were energy minimized to allow for relaxation of potential clashes or energetically disfavorable rotamers and solvent or ion interactions. (*B*) The simulations ran for 1000 ns and then checked for equilibration by calculation of the root mean square displacement (RMSD) in comparison to the structure after minimization and equilibration. Data before 600 ns was cleaved from the analysis datasets.

**Table S1.** Quantification of nucleosomal dimensions by AFM analysis. Data demonstrate that *in vitro* chromatin reconstitution yields equivalent dimensions for CENP-A and H3, butthat CENP-A nucleosomes increase in height by ∼0.4nm when bound to CENP-CCD. Heights (nm), Diameters (nm), and volumes (nm3) were calculated for representative particles of each class of nucleosome imaged by atomic force microscopy in-fluid conditions (*Methods*).

| Supplemental Table S1: Quantification of nucleosome particles in H3, CENP-A, and CENP-A + CENP-C<sup>CD</sup> conditions. |  |  |  |  |  |  |  |  |  |  |  |
| --- | --- | --- | --- | --- | --- | --- | --- | --- | --- | --- | --- |
| H3 |  |  | CENP-A |  |  | 2X CENP-C<sup>CD</sup> |  |  | 4X CENP-C<sup>CD</sup> |  |  |
| height(nm) | diameter (nm) | volume(nm<sup>3</sup>) | height(nm) | diameter (nm) | volume(nm<sup>3</sup>) | height(nm) | diameter (nm) | volume(nm<sup>3</sup>) | height(nm) | diameter (nm) | volume(nm<sup>3</sup>) |
| 3.8 | 12.3 | 300.8 | 3.7 | 12.8 | 317.2 | 5.4 | 12.5 | 441.5 | 4.7 | 14.1 | 489.1 |
| 4.1 | 12.3 | 324.6 | 3.7 | 13.4 | 347.6 | 4.5 | 15.1 | 536.9 | 3.6 | 15.1 | 429.5 |
| 3.2 | 15.1 | 381.8 | 3.1 | 12.5 | 253.4 | 4.4 | 12.8 | 377.2 | 5.4 | 12.8 | 463.1 |
| 3.5 | 16.7 | 510.8 | 3.2 | 14.4 | 347.2 | 4.3 | 12.8 | 368.6 | 4.4 | 14.2 | 464.4 |
| 3.7 | 13.5 | 352.8 | 3.2 | 12.6 | 265.8 | 4.3 | 14.1 | 447.3 | 3.7 | 14.8 | 424.1 |
| 3.9 | 14.3 | 417.3 | 3.5 | 13.9 | 353.8 | 4.3 | 12.7 | 362.9 | 3.6 | 12.5 | 294.3 |
| 4.2 | 13.5 | 400.5 | 3.5 | 13.5 | 333.8 | 4.2 | 13.4 | 394.6 | 3.6 | 12.7 | 303.7 |
| 3.9 | 15.5 | 490.3 | 3.6 | 15 | 423.9 | 4.1 | 14.5 | 451.1 | 3.3 | 15.1 | 393.7 |
| 3.9 | 14.3 | 417.3 | 3.6 | 13.8 | 358.7 | 4 | 13.1 | 359.2 | 3.8 | 13.6 | 367.8 |
| 4 | 12.5 | 327.1 | 3.8 | 14.2 | 400.9 | 3.9 | 13.5 | 371.9 | 4.5 | 15.9 | 595.4 |
| 3.7 | 15.5 | 465.2 | 3.9 | 14 | 400.1 | 3.8 | 12.8 | 325.8 | 4.1 | 14.7 | 463.5 |
| 3.7 | 12.7 | 312.3 | 4.1 | 14.1 | 426.5 | 3.8 | 13.6 | 367.8 | 3.8 | 14.1 | 395.6 |
| 3.7 | 13.5 | 352.8 | 4.5 | 13.6 | 435.5 | 4.8 | 11.9 | 355.7 | 3.7 | 17.5 | 593.1 |
| 4 | 12.3 | 316.7 | 4.4 | 12.8 | 377.2 | 4.2 | 13.8 | 418.5 | 3.3 | 15.5 | 414.9 |
| 3.9 | 14.3 | 417.3 | 4.2 | 12.6 | 348.9 | 4.1 | 13.5 | 391.1 | 5.1 | 15.1 | 608.5 |
| 4 | 12.5 | 327.1 | 4 | 12.8 | 342.9 | 3.7 | 14.1 | 384.9 | 4.4 | 16.7 | 642.1 |
| 3.7 | 15.5 | 465.2 | 4 | 14.6 | 446.2 | 3.4 | 12.8 | 291.5 | 4.4 | 12.5 | 359.7 |
| 3.3 | 15.5 | 414.9 | 3.9 | 15 | 459.2 | 3.3 | 15.1 | 393.7 | 4.2 | 13.5 | 400.8 |
| 3.8 | 14.3 | 406.6 | 3.9 | 13.9 | 394.3 | 3.7 | 13.5 | 352.8 | 4.1 | 14.2 | 432.5 |
| 4.1 | 14.3 | 438.7 | 3.8 | 13.8 | 378.7 | 3.9 | 12.9 | 339.6 | 3.9 | 12.6 | 324.1 |
| 3.7 | 15.5 | 465.2 | 3.7 | 14.5 | 407.1 | 4.4 | 12.9 | 383.1 | 3.8 | 13.8 | 378.7 |
| 4.6 | 15.5 | 578.3 | 3.5 | 12.5 | 286.1 | 3.6 | 13.2 | 328.2 | 3.7 | 14.1 | 384.6 |
| 4 | 13.1 | 359.2 | 3.4 | 12.8 | 291.5 | 4.1 | 13.6 | 396.8 | 5.6 | 11.9 | 415.1 |
| 4.4 | 12.7 | 371.3 | 3.3 | 14.2 | 348.2 | 4.2 | 13.9 | 424.6 | 5.4 | 14.3 | 577.8 |
| 3.4 | 13.9 | 343.7 | 4.1 | 13.6 | 396.8 | 4.2 | 15 | 494.5 | 4.4 | 13.9 | 444.8 |
| 3.6 | 12.7 | 303.8 | 4.1 | 14.5 | 451.1 | 4.3 | 13.7 | 422.3 | 4.3 | 13.1 | 386.1 |
| 4.1 | 13.9 | 414.5 | 4 | 12.5 | 327.1 | 4.6 | 14.9 | 534.4 | 4.2 | 11.9 | 311.2 |
| 4.1 | 13.1 | 368.2 | 3.8 | 11.9 | 281.6 | 4.7 | 12.3 | 372.1 | 4.1 | 13.1 | 368.2 |
| 4.3 | 13.9 | 434.7 | 3.8 | 14.9 | 441.5 | 4.9 | 14.1 | 509.8 | 3.9 | 13.9 | 394.3 |
| 3.8 | 14.3 | 406.6 | 3.8 | 13.2 | 346.5 | 4.5 | 12.7 | 379.8 | 5.4 | 13.2 | 492.4 |
| 3.6 | 11.9 | 266.7 | 3.7 | 13.6 | 358.1 | 4.5 | 12.8 | 385.8 | 4.5 | 12.3 | 356.2 |
| 3.8 | 11.9 | 281.6 | 3.7 | 12.8 | 317.2 | 4.5 | 14.2 | 474.8 | 4.1 | 13.2 | 373.8 |
| 3.6 | 13.1 | 323.3 | 3.7 | 12.9 | 322.2 | 4.4 | 13.2 | 401.2 | 4.1 | 14.5 | 451.1 |
| 4.1 | 13.9 | 414.5 | 3.7 | 15.6 | 471.2 | 4.1 | 14.2 | 432.6 | 3.9 | 15 | 459.2 |
| 3.8 | 13.9 | 384.2 | 4.1 | 15.2 | 495.7 | 4.1 | 13.2 | 373.8 | 3.7 | 14.2 | 390.4 |
| 4.1 | 15.1 | 489.2 | 3.7 | 12.9 | 322.2 | 3.9 | 14.5 | 429.1 | 3.5 | 13.9 | 353.8 |
| 4.1 | 14.7 | 463.6 | 3.6 | 13.7 | 353.6 | 3.8 | 12.5 | 310.7 | 3.5 | 14.6 | 390.4 |
| 3.5 | 12.1 | 268.1 | 3.6 | 13.6 | 348.4 | 3.7 | 12.9 | 322.2 | 3.4 | 14.1 | 353.7 |
| 3.4 | 13.9 | 343.7 | 3.6 | 16.2 | 494.4 | 3.4 | 14.7 | 384.4 | 3.5 | 13.7 | 343.7 |
| 3.7 | 14.3 | 395.9 | 3.5 | 14.6 | 390.4 | 3.2 | 12.6 | 265.8 | 3.6 | 14.7 | 407.1 |
| 3.4 | 15.5 | 427.4 | 3.4 | 12.7 | 286.9 | 3.6 | 12.7 | 303.8 | 3.8 | 12.9 | 330.9 |
| 4.1 | 13.9 | 414.5 | 3.4 | 15.1 | 405.7 | 3.7 | 14.3 | 395.9 | 3.9 | 14.6 | 435.1 |
| 3.8 | 14.7 | 429.7 | 3.4 | 14.4 | 368.9 | 4.1 | 14.7 | 463.6 | 4.1 | 13.9 | 414.5 |
| 3.7 | 15.9 | 489.5 |  |  |  | 4.3 | 14.2 | 453.7 | 4.2 | 14.1 | 436.9 |
| 3.5 | 14.7 | 395.8 |  |  |  | 4.5 | 12.1 | 344.7 | 4.5 | 13.8 | 448.4 |
|  |  |  |  |  |  |  |  |  | 4.9 | 14.6 | 546.6 |
|  |  |  |  |  |  |  |  |  | 5.9 | 12.7 | 498.1 |

**Table S2.** Quantification of chromatin folding demonstrates that CENP-C increases CENP-A chromatin clustering. Native chromatin incubated with or without the CENP-C fragment was visually inspected on AFM and identified as “open” or “clustered”. Total number of CENP-A clusters/total number of CENP-A nucleosome arrays per scan. Both analyses demonstrate that CENP-<u>C incre</u>ases CENP-A chromatin clustering (*Methods*).

|  | ACA WT |  |  | ACA CENP-C<sup>CD</sup> |  |  |
| --- | --- | --- | --- | --- | --- | --- |
|  | # arrays | # clsuters | # clusters/# arrays | # arrays | # clsuters | # clusters/# arrays |
| **Scan1** | 14 | 5 | 0.36 | 19 | 7 | 0.37 |
| **Scan2** | 19 | 6 | 0.32 | 13 | 5 | 0.38 |
| **Scan3** | 22 | 6 | 0.27 | 14 | 6 | 0.43 |
| **Scan4** | 17 | 7 | 0.41 | 26 | 12 | 0.46 |
| **Scan5** | 34 | 13 | 0.38 | 15 | 6 | 0.40 |
| **Scan6** | 48 | 13 | 0.27 | 22 | 9 | 0.41 |
| **Scan7** | 40 | 16 | 0.40 | 24 | 11 | 0.46 |
|  | ACA WT |  |  | ACA CENP-C<sup>CD</sup> |  |  |
|  | # arrays | # clsuters | # clusters/# arrays | # arrays | # clsuters | # clusters/# arrays |
| **Scan1** | 31 | 10 | 0.32 | 38 | 28 | 0.74 |
| **Scan2** | 34 | 11 | 0.32 | 50 | 27 | 0.54 |
| **Scan3** | 26 | 14 | 0.54 | 49 | 35 | 0.71 |
| **Scan4** | 19 | 7 | 0.37 | 55 | 39 | 0.71 |
| **Scan5** | 28 | 8 | 0.29 | 43 | 21 | 0.49 |

**Table S3.** RNAP2 levels on centromeric chromatin are reduced under CENP-C over-expression conditions. Cells were transfected (or not) with full length CENP-C which was over-expressed (OE) for 3 days, native centromeric chromatin was extracted by CENP-C or ACA ChIP, from wildtype cells or CENP-C OE cells, in parallel. Chromatin was evaluated for RNAP2and CENP-A occupancy on Western blots. 3 independent replicates were quantified using the Licor scanner and automated software. Quantification of RNAP2 in CENP-C IP or ACA IP over Input demonstrates a suppression of RNAP2 levels on centromeric chromatin upon CENP-C OE, and a reduction of total CENP-A levels when RNAP2 is diminished.

| Supplemental Table S3: Quantification of band intensity of RNA polymerase 2 and CENP-A western blots. Values are arbitrary units derived from LiCor's software (10<sup>3</sup>). |  |  |  |  |  |  |  |  |
| --- | --- | --- | --- | --- | --- | --- | --- | --- |
| RNAP2 |  |  |  |  |  |  |  |  |
|  | WT |  |  |  |  | CENP-C<sup>OE</sup> |  |  |
|  | Input | CENP-C | ACA |  |  | Input | CENP-C | ACA |
| Exp1 | 5.92 | 26.8 | 17.5 |  | Exp1 | 6.52 | 9.12 | 8.76 |
| Exp2 | 7.76 | 14.1 | 12.6 |  | Exp2 | 9.36 | 7.5 | 10.8 |
| Exp3 | 7.34 | 24.9 | 17.8 |  | Exp3 | 6.31 | 8.07 | 7.67 |
|  |  | Ratio CpC/input | Ratio ACA/input |  |  |  | Ratio CpC/input | Ratio ACA/input |
| Exp1 |  | 4.53 | 2.96 |  | Exp1 |  | 1.40 | 1.34 |
| Exp2 |  | 1.82 | 1.62 |  | Exp2 |  | 0.80 | 1.15 |
| Exp3 |  | 3.39 | 2.43 |  | Exp3 |  | 1.28 | 1.22 |
|  | mean | 3.25 | 2.33 |  |  | mean | 1.16 | 1.24 |
|  | StDev | 1.36 | 0.67 |  |  | StDev | 0.32 | 0.10 |
| CENP-A |  |  |  |  |  |  |  |  |
|  | WT |  |  |  |  | CENP-C<sup>OE</sup> |  |  |
|  | Input | CENP-C | ACA |  |  | Input | CENP-C | ACA |
| Exp1 | 0.98 | 8.63 | 13.2 |  | Exp1 | 1.97 | 6.74 | 8.53 |
| Exp2 | 0.57 | 10 | 11.4 |  | Exp2 | 0.56 | 8.7 | 2.18 |
| Exp3 | 0.38 | 1.71 | 2.23 |  | Exp3 | 0.52 | 1.6 | 2.02 |
|  |  | Ratio CpC/input | Ratio ACA/input |  |  |  | Ratio CpC/input | Ratio ACA/input |
| Exp1 |  | 8.81 | 13.47 |  | Exp1 |  | 3.42 | 4.33 |
| Exp2 |  | 17.54 | 20.00 |  | Exp2 |  | 15.54 | 3.89 |
| Exp3 |  | 4.50 | 5.87 |  | Exp3 |  | 3.08 | 3.88 |
|  | mean | 10.28 | 13.11 |  |  | mean | 7.34 | 4.04 |
|  | StDev | 6.65 | 7.07 |  |  | StDev | 7.10 | 0.25 |

## Notes

#### Summary of Updates

The original manuscript, first posted in September 2017, was split into two in July 2018, to highlight the two major findings separately. The second manuscript will be posted on biorXiv shortly.

